# The role of endospore appendages in spore-spore contacts in pathogenic bacilli

**DOI:** 10.1101/2024.04.22.590507

**Authors:** Unni Lise Jonsmoen, Dmitry Malyshev, Mike Sleutel, Elise Egeli Kristensen, Ephrem Debebe Zegeye, Han Remaut, Magnus Andersson, Marina Elisabeth Aspholm

## Abstract

Species within the spore-forming *Bacillus cereus* sensu lato group are recognized for their role in food spoilage and food poisoning. *B. cereus* spores are decorated with numerous pilus-like appendages, called S-ENAs and L-ENAs. These appendages are believed to play crucial roles in self-aggregation, adhesion, and biofilm formation. By using both bulk and single-cell approaches, we investigate the role of S-and L-ENAs as well as the impact of different environmental factors in spore-to-spore contacts and in the interaction between spores and vegetative cells. Our findings reveal that ENAs, and particularly their tip fibrilla, play an essential role in spore self-aggregation but not in the adhesion of spores to vegetative cells. The absence of L-BclA, which builds the L-ENA tip fibrillum, reduced both S-and L-ENA mediated spore aggregation, emphasizing the interconnected roles of S-and L-ENAs. Increased salt concentrations in the liquid environment significantly reduced spore aggregation, implying a charge dependency of spore-spore interactions. By elucidating these complex interactions, our study provides valuable insights into spore dynamics. This knowledge can guide future studies on spore behavior in environmental settings and aids in developing strategies to manage bacterial aggregation for beneficial purposes, like controlling biofilms in food production equipment.

## Introduction

Members of *Bacillus cereus sensu lato* (*s.l.*) group (*B. cereus* spp.) are widespread in various natural environments, including air, water, vegetation, and soil. These facultative anaerobic Gram-positive bacteria are frequent contaminants of raw food material (Rahnama et al., 2023) and food production equipment, leading to both emetic and diarrheal types of food poisoning (Shaheen et al., 2010a; Yang et al., 2023). An additional concern is the release of various enzymes that degrade food components, resulting in decreased food quality and shortened shelf life. A major challenge for food producers is the ability of *B. cereus* spp. to form highly resilient endospores (spores), which makes the decontamination of food products and production equipment a difficult and costly task. These spores can withstand harsh conditions such as wet heat, irradiation, chemicals, and desiccation — conditions that would normally eliminate the vegetative cells. The endospores’ resilience is attributed to several factors, including a dehydrated core enriched with calcium-dipicolinic acid (CaDPA) and protective small acid-soluble proteins, alongside several semi-permeable layers, with each layer providing unique protection (Setlow, 2006; Sunde et al., 2009; Swick et al., 2016).

It has been shown that the *B. cereus* spores are more adhesive than their vegetative counterpart and bind more readily to materials found in food production facilities, such as stainless steel, plastic, and glass surfaces (Peng et al., 2001; Rönner et al., 1990). The high adhesiveness complicates their removal (Faille et al., 2016) and provides increased resistance to environmental challenges (Simmonds et al., 2003). Furthermore, adhered spores may serve as a foundation for the development of biofilm on these surfaces (Faille et al., 2014). Biofilms attach to surfaces and establish a dynamic microenvironment that promotes collective survival and adaptation (Kragh et al., 2016). Consequently, the presence of biofilms in food production equipment poses a persistent contamination risk, exhibiting strong resistance to removal and resulting in substantial costs related to cleaning and sanitation within the food industry. To improve the efficiency of spore removal methods, additional insights into the interactions between spores and their surroundings are needed. Bacterias’ ability to interact and autoaggregate (adhere to one another) also contributes to their resilience (Burel et al., 2021). Autoaggregation is common among environmental and clinical bacterial strains and is associated with protection against environmental stresses and immune responses (Demirdjian et al., 2019). The process of bacterial autoaggregation is typically mediated by specific adhesion molecules on the bacterial surface, such as proteins, polysaccharides, and longer surface structures (Dogsa et al., 2023). Several studies have highlighted the role of structures like pili, including chaperone-usher pili (Faris et al., 1983), type IV pili (Bieber et al., 1998) and curli (Boyer et al., 2007), in facilitating this self-adhesion in bacteria such as *Yersina eneterocolitica* and *Escherichia coli*. Whereas the molecular determinants of autoaggregation of vegetative cells are well described, little is known about the aggregation process seen in spores.

Recent studies have revealed that most *B. cereus* spp. carry genes for pilus-like structures known as spore-specific endospore appendages (ENAs) (Pradhan et al., 2021). The architecture of two types of ENA, denoted S-and L-ENA, expressed on the surface of a food-borne outbreak isolate of *Bacillus paranthracis* NVH 0075/95, was recently solved (Pradhan et al., 2021; Sleutel et al., 2023). The S-ENAs are described as “staggered”, μm long proteinaceous fibers that are tens of nm wide. They end in “ruffles” featuring four to five thin tip fibrils that in general appearance resemble the tip fibrils found on adhesive pili like type 1 pili and P-pili of uropathogenic *E. coli* (Pradhan et al., 2021). The S-ENAs, which are the most abundant type of appendages on NVH 0075/95 spores, are believed to be anchored beneath the exosporium layer, directly to the spore coat. In contrast, the “ladder-like” L-ENAs, are thinner and shorter than the S-ENAs and are linked to the exosporium layer via the ExsL protein (formerly denoted CotZ) (Sleutel et al., 2023). Each L-ENA fiber is decorated with a single tip fibrillum composed of the collagen-like L-BclA protein (Sleutel et al., 2023). Notably, deleting the *l-bclA* gene leads to L-ENAs lacking the tip fibrillum, while it does not affect the presence of tip fibrilla on S-ENAs’. This distinction in tip fibril and the structural differences between the two types of ENAs suggests specific roles in *B. cereus* spp. spores, as highlighted in recent research (Sleutel et al., 2023).

Until recently, the lack of structural and genetic data on ENAs has limited understanding of their function in *B. cereus* spp. As reviewed in Zegeye et al., these fibers may play a role in adherence to surfaces and biofilm formation (Zegeye et al., 2021). In this context, it is important to note that a mono-species biofilm of *B. cereus* spp. can comprise as much as 90% spores (Wijman et al., 2007), suggesting that the ENAs may constitute a substantial part of the biofilm matrix. Further research into the S-ENA revealed that they are rigid yet flexible fibers, maintaining integrity under extension, which supports their proposed role in facilitating spore-to-spore binding and potentially enhancing autoaggregation (Jonsmoen et al., 2023). The limited understanding of autoaggregation in *B. cereus* spp. spores prompts the need for a focused investigation, driven by the potential implications of this phenomenon on biofilm formation, environmental persistence, and, ultimately, food safety.

In this work, we examine the role of S-ENA and L-ENA, as well as the tip fibrillum of L-ENAs, in the self-aggregation behavior of *B. cereus* spore populations, using both wildtype (WT) and ENA-deficient mutant spores (Fig 1A). We investigate their roles in spore-spore and spore-vegetative cell interactions, focusing on how these interactions are influenced by factors such as ionic strength and surfactants in the surrounding liquid. Our findings provide insights into the mechanisms that potentially enhance spore resistance to environmental stress and aid in biofilm formation. These insights could contribute to the development of more efficient strategies for removing *B. cereus* spores from food production environments and equipment. Accordingly, it has the potential to not only save costs and reduce food waste but also promote more sustainable dairy production practices.

**Fig 1.**
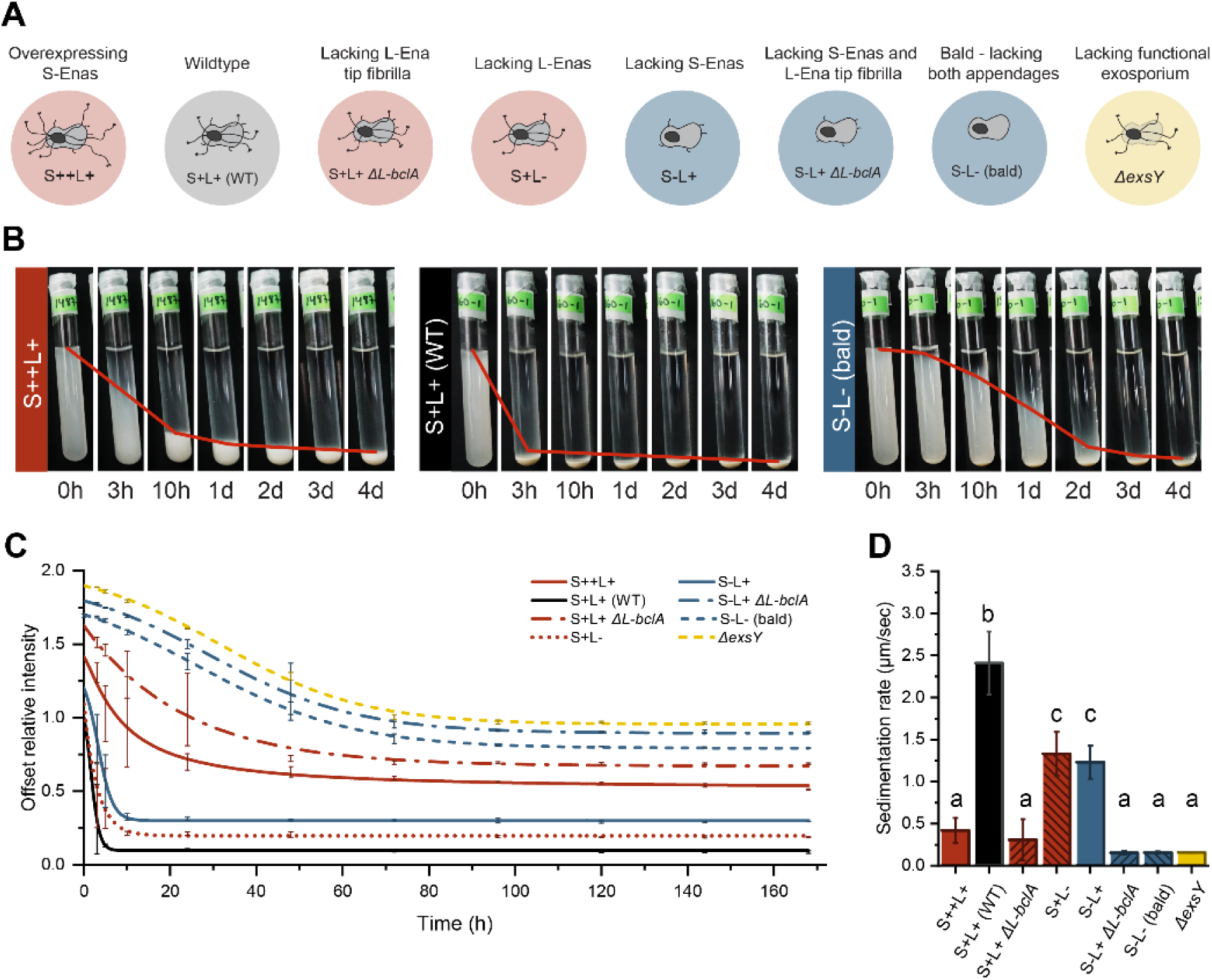
Sedimentation dynamics. Schematic overview of the strains used in the study (A). Timelapses showing spores of three strains sedimenting in a glass tube over a period of four days (B): the S++L+ strain (left), WT spores (middle), and bald spores (right). The red line represents the 50% intensity mark for the corresponding sample. Sedimentation patterns of WT spores and all seven mutants in a water suspension, fitted with Boltzmann curves (C) For overlaying sedimentation patterns see Figure S1. Sedimentation rate is displayed in µm/sec (D). Bars that share a letter are not significantly different, as indicated by Tukey’s grouping (p<0.05). Numerical averages are given in Table S2.

## Experimental procedures

### Strains and spore preparations

The food outbreak isolate *B. paranthracis* NVH 0075/95 (Lund & Granum, 1996) and the isogenic mutants used in this paper are listed in Table 1 and their phenotype illustrated in Fig 1. For preparation of spores, the bacteria were streaked onto LB agar plates and incubated for no less than two weeks at 37°C. When approximately 98% of the bacteria had sporulated, as determined by phase contrast microscopy (Olympus BX51, Olympus Corporation, Japan), the spores were harvested as described before in Jonsmoen et al (2023) (Jonsmoen et al., 2023). Pure spore suspensions were stored at 4 °C until the start of the experiment.

**Table 1.** List of *Bacillus paranthracis* NVH 0075/95 strains included in this study.

| Name | Genotype | S-ENA | L-ENA | L-BclA | Morphology | Reference |
| --- | --- | --- | --- | --- | --- | --- |
| S+L+ (WT) | WT | + | + | + | Wild type. Both S-ENAs and L-ENAs. | (Lund & Granum, 1996) |
| S+L- | *Δena3* | + | - |  | No L-ENA fibers, but S-ENAs are present | (Jonsmoen et al., 2023; Sleutel et al., 2023) |
| S-L+ | *Δena1ABC* | - | + | + | No S-ENA fibers, but L-ENAs present | (Pradhan et al., 2021) |
| S-L- (bald) | *Δena1ABC Δena3* | - | - |  | Bald spores, neither S nor L-ENA present | (Jonsmoen et al., 2023; Sleutel et al., 2023) |
| *ΔexsY* | *ΔexsY* | + | - |  | S-ENA present, but no L-ENA | (Jonsmoen et al., 2023) |
| S++L+ | *Δena1B::pHT304-ena1AB* | ++ | + |  | More and longer S-ENA are expressed. L-ENAs are present. | (Jonsmoen et al., 2023; Pradhan et al., 2021) |
| *ΔL-bclA* | *Δl-bclA* | + | + | - | No L-ENA tip fibrilla | (Sleutel et al., 2023) |
| S-L+ *ΔL-bclA* | *Δena1ABC Δl-bclA* | - | + | - | No S-Ena. The L-ENA stem is present, but lacking tip-fibrilla. | This study |

### Sedimentation assays

To study the sedimentation behavior, three batches of spores, suspended in sterile distilled water, were pooled and the OD_600_ of the spore suspensions were adjusted to 10, before being mixed thoroughly by vortexing for about 15 sec. The suspensions were placed in three borosilicate sample tubes (14 mm X 130 mm, *DWK Life Sciences*) and left to settle. A SONY ILCE-5100 camera with a SELP1650 lens was used to capture 24.3 megapixel images of the tubes at 0, 3, 5, and 10 hours. Subsequently, images were taken every 24 hours until the samples had fully settled.

To verify that the spores had not germinated during the experiment, all samples were homogenized after the experiment had ended and spore status were inspected using phase contrast microscopy (Olympus BX51, Olympus Corporation, Japan).

### Optical tweezer setup

For the analysis of binding between single spores or between a spore and a single vegetative cell, we used an optical tweezer (OT) setup integrated in an inverted microscope (Olympus IX71, Olympus Corporation, Japan) integrated with a water immersion objective (UPlanSApo60XWIR, 60×/1.2 N.A, Olympus) (Stangner et al., 2018). The samples were recorded in bright field mode using a 1920 × 1440 pixel complementary metal oxide semiconductor (CMOS) camera (C11440-10C, Hamamatsu Photonics, Japan). The OT system has been optimized to reduce fluctuations using the Allan variance analysis as described in (Andersson et al., 2011) placed on the sample holder, which is temperature-controlled and maintained at 25°C. A 1064 nm DPSS laser (Rumba, 05-01 Series, Cobolt AB, Sweden) was used for trapping spores and bacterial cells. We made sure that the irradiation dose was below the threshold for disrupting the spore bodies, about 50 J, when trapping the spores (Malyshev et al., 2022).

### OT catch-and-release test

Three types of catch-and-release tests were conducted to observe the role of ENAs in adherence. The first analysis was done on isogenic spores, where 1 µl of concentrated spore suspension was added to the middle of a sample slide (24 × 60 mm, no. 1, Paul Marienfeld, Lauda-Königshofen, Germany) with double sticky tape (product no. 34-8509-3289-7, 3M) on both sides. Then, 10 µl of the suspension media, either Milli-Q water, PBS (137 mM NaCl, 1.7 mM KCl, 10 mM Na_2_HPO_4_, 1.8 mM KH_2_PO_4_) at different concentration (1:1, 1:2, 1:10), 0.05% Tween-20 (BP337-100, Fisher BioReagents™) or 1% BSA (A6003, Sigma-Aldrich), was added on top of the concentrated spore suspension. The sample was then enclosed with a coverslip (20 × 20 mm, no. 1, Paul Marienfeld), and the edges sealed off. Two spores were captured in the trap of the optical tweezer and brought to the spore-free zone closer to the edges of the sample and held together for approximately 5 seconds in the optical trap using a holding a power of 1 W and a 1064 nm laser. After the spores were released from the trap, we followed them for at least 1 minute, to see if the spores stayed attached to each other or drifted apart by Brownian motion. The interaction process was recorded, and the outcome was noted. A total of 30 interactions were initiated for each experiment, except for the WT strain (S+L+) where 60 interactions were initiated.

The second catch-and-release test was done on vegetative cells or on spores of different genotypes. A channel was made by adding a double sticky tape on both sides of the sample to create a channel, closed off by a coverslip. A volume of 2 µl of each spore suspension to be tested was applied at either end of the channel so that a spore-free zone was created in the middle. Bald spores were deposited at one side of the channel, while another spore type was deposited on the opposite side (illustrated in Fig. 6). The channel was then filled with Milli-Q water. One spore from each side of the channel was then brought to the spore-free middle zone by using the OT. As in the prior experiment, the spores were held together, released and the resulting outcome was noted.

For catch and release analysis of interaction between a single spore and a vegetative cell, strain NVH 0075/95 was grown overnight in LB at 30°C under shaking at 200 rpm. The following day, the overnight culture was diluted 1:100 with fresh medium before left to grow for 3 hours. The bacteria were pelleted by centrifugation at 2000 rpm for 5 minutes. The supernatant was discarded, and the pellet was resuspended in Milli-Q water. The catch-and-release test was performed following the same procedure used for testing binding between individual spores, except that in this case, spores were added on one side and vegetative bacteria on the other.

### Electron microscopy sample preparation

To obtain images of vegetative cells, 100 µL of an LB overnight culture of strain NVH 0075/95 was transferred to a fresh media and left to grow for 4 hours at 37°C under shaking at 200 rpm. The cells were then centrifugated down for 10 minutes at low speed (2500 rpm), and the pellet was washed once in PBS before fixated (2% paraformaldehyde). A copper grid (400 mesh) covered with FCF400-CU Formvar Carbon Film was placed on top of a droplet of the fixed cell suspension for 1 minute. The excess suspension was removed by capillary force using dry filter paper. The grid was transferred to a droplet of 4% uranyl acetate staining for another minute before the dried grid was ready for analysis using a JEM-2100Plus Electron Microscope (JEOL Ltd., Japan).

### S-ruffle measurements

For the S-ENA ruffle measurements, we first collected raw negative stain micrographs. For this, a suspension of spores was applied onto formvar/carbon-coated copper grids (Electron Microscopy Sciences) with a 400-hole mesh. The grids were glow-discharged (ELMO; Agar Scientific) with 4mA plasma current for 45 seconds. 3 μl of a bacterial spore suspension was applied onto the glow-discharged grids and left to adsorb for 1 minute. The solution was dry blotted, followed by three washes with 15 μl Milli-Q. Next, 15 μl drops of 2% uranyl acetate were applied three times for 10 seconds, 2 seconds, and 1 minute respectively, with a blotting step in between each application. The excess uranyl acetate was then dry blotted with Whatman type 1 paper. All grids were screened with a 120 kV JEOL 1400 microscope equipped with LaB6 filament and TVIPS F416 CCD camera. Next, micrographs were loaded into ImageJ 1.54f and scaled using a pixel resolution of 1.94Å/pix. Ruffle lengths were measured using the segment tool by drawing a segmented line starting from the tip of the S-ENA pilus to the terminus of the tip fibrillum, following the curvature of the ruffle.

### Data analysis and statistics

Images from the sedimentation assays were analyzed using ImageJ 1.54f (Rueden et al., 2017). The size of the pellet after 6 days was determined by measuring it against an internal standard on the images. Additionally, the pixel intensity of the spore suspensions was assessed by generating a profile plot, normalizing it according to the highest value, and then balancing it into 200 datapoints using a randomized algorithm in RStudio (Version 2023.12.1). The intensity profiles were plotted against time and datapoints crossing 50% light intensity were extracted and plotted as a function of time. ImageJ was also used to estimate the center-to-center distances (µm) between adhered spores or spores adhered to vegetative cells using the line tool. Each interaction was measured three times during the recording, and the average was noted.

Using linear regression analysis on the linear phase of the settling process, the spores’ sedimentation rate was estimated. Further, Tukey’s test in RStudio was applied to determine significant differences between the WT and mutant strains. Graphs were plotted in Origin 2024 (OriginLabs, Version 10.1.0.178). We used the non-linear curve fitting tool in Origin to fit the segmentation data to Boltzmann-type curves, aiding visual interpretation.

## Results

### S-and L-ENA influence spore sedimentation

The interaction among bacterial cells or spores within a population is often assessed by observing their sedimentation patterns in liquid medium (Nwoko & Okeke, 2021; Trunk et al., 2018). This technique takes advantage of the modified sedimentation behavior arising from cell-cell adhesion and the formation of aggregates. Cells or spores that closely aggregate or form clumps will sediment at a faster rate than individual cells because the combined mass of the aggregate is greater, leading to increased gravitational settling. To investigate the role of appendages in spore aggregation, we utilized this sedimentation approach and set up a series of glass tubes containing a spore suspension and left them to settle over one week. Illustrative data for three spore populations, captured at regular time intervals to show the sedimentation patterns of the spore suspensions, are compiled and presented in Fig 1B. Depending on the appendage combination expressed on the spores, the spore suspensions showed different sedimentation behavior. The WT spores, which express both S-and L-ENAs, showed the fastest sedimentation rate, resulting in the rapid formation of a dense pellet at the bottom of the test tubes. The bald spores, which lack both S-and L-ENAs, showed a much slower and gradual sedimentation behavior than WT spores. Spores of the S++L+ strain, which expresses ∼3x longer S-ENAs compared WT spores (Pradhan et al., 2021), sedimented slower than WT spores and formed a less dense pellet at the bottom of the tube.

Based on their sedimentation pattern, all tested spore populations (Fig 1A) could be categorized into the fast or slow group as shown in (Fig 1B). The first group comprises spores of the WT strain, S+L-spores (lacking L-ENAs) and S-L+ spores (lacking S-ENAs), all of which show a similar and fast sedimentation behavior. This suggests that these spore groups form aggregates that have a lower surface area relative to their volume compared to individual cells. Thus, since the drag force acts on the surface of the cells, a lower surface area to volume ratio results in less resistance against sedimentation. The second group consists of bald spores, the Δ*exsY* spores that express a deficient exosporium and L-BclA depleted spores that lack tip fibrillum on L-ENA (Sleutel et al., 2023). To determine the sedimentation rate, we used image analysis to identify the position in the column where the pixel intensity reached 50% of its maximum value. Additionally, we calculated the sedimentation rate of each strain by conducting a regression analysis on the linear segment of the sedimentation curve, Fig 1D.

After six days (144 h), most of the spore suspensions had settled at the bottom of the tubes and it became clear that the size reflecting the density of the pellet was affected by the ENA phenotype and the presence of the exosporium (Fig 2A). The Δ*exsY* strain, which lacks an exosporium, formed the smallest and densest pellet with an average size of 1.0 ± 0.1 mm, while the largest and least dense pellets were formed by the strain overexpressing S-ENA (S++L+), with an average pellet size of 3.6 ± 0.1 mm. The absence of L-BclA appeared to impact pellet density; spore populations lacking L-BclA (S+L+ Δ*L-bclA* and S-L+ Δ*L-bclA*) displayed smaller pellet sizes compared to their counterparts that retained the intact tip fibrillum on the L-ENA fiber.

**Fig 2.**
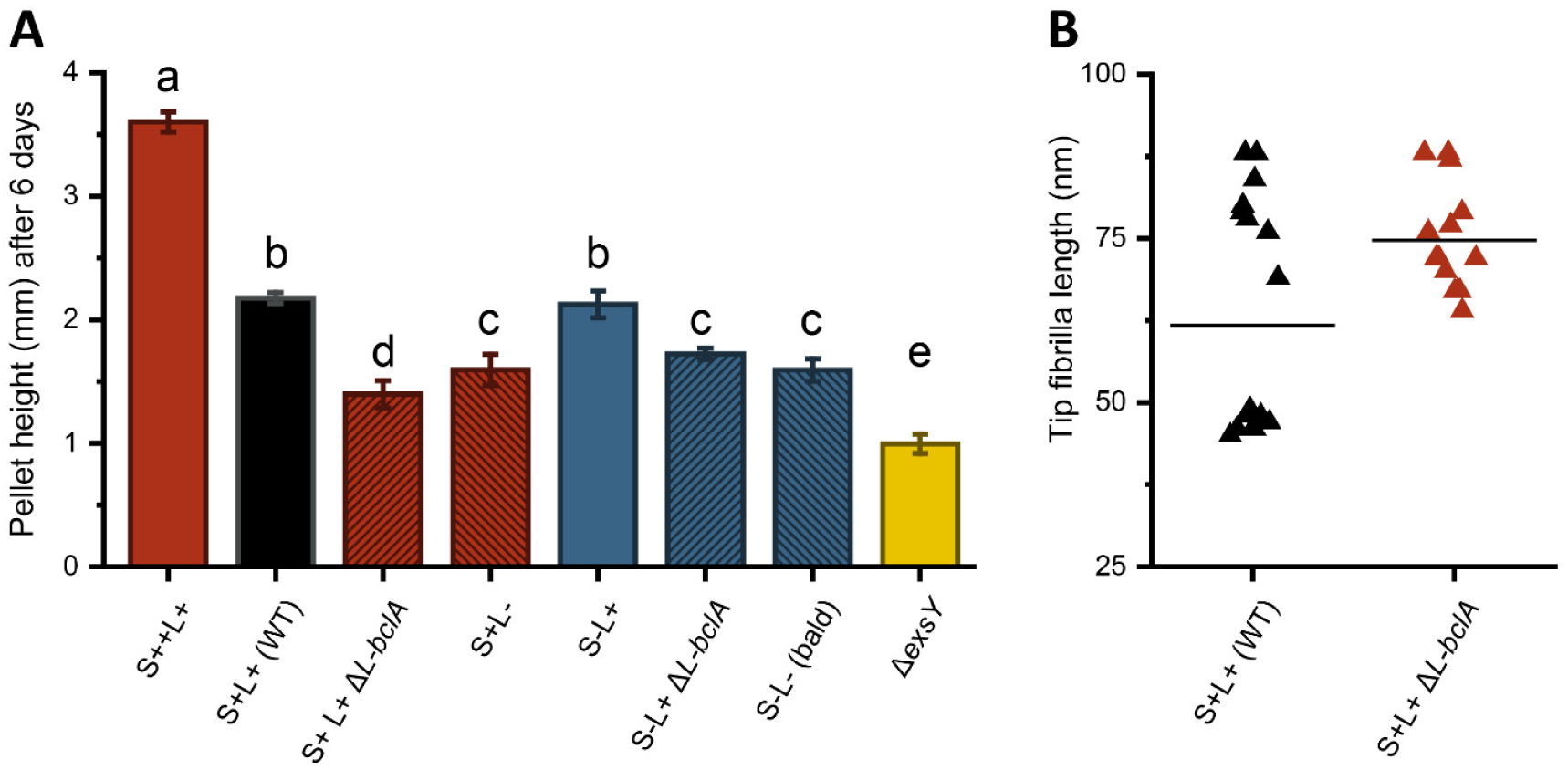
Pellet size and S-ENA tip fibrilla distributions. Pellet size after six days (144 h). Bars that share a letter are not significantly different, as indicated by Tukey’s grouping (p<0.05). Numerical averages are given in Table S2(A). Length distribution of S-ENA tip fibrilla on S+L+ (WT) and S+L+ Δ*L-bclA* (B) from TEM images. The line represents the mean fibrilla length.

Because of the observed decrease in sedimentation rate for L-BclA deficient spores (S+ L+ Δ*L-bclA)* compared to those of the S+ L-strain, we aimed to investigate whether the depletion of *L-bclA* impacts the tip fibrilla distribution on the S-ENAs. Upon analyzing TEM images of S-ENA on WT (S+ L+) and L-BclA deficient spores, we observed no difference in the number of tip fibrilla per appendage fiber (Sleutel et al., 2023). However, when we measured the length of the individual S-ENA fibrils in TEM images for both WT (S+ L+) and S+L+ Δ*L-bclA*, we detected a variation in the length distribution of the tip fibrilla fibers. From Fig 2B, we see a cluster of shorter tip fibrilla in the WT, which is lacking in the S+L+ Δ*L-bclA* measurements. These shorter tip fibrilla are of similar length as the L-BclA, 47 ± 1 nm and 45 nm (Sleutel et al., 2023), respectively.

### Appendages facilitate close interactions among spores

To further test the role of ENAs in spore-spore interactions, we performed a catch-and-release analysis of spore-to-spore binding using an optical tweezers (OT) system. This allowed us to capture two spores and keep them in close proximity for a set duration (Fig 3a-c). Upon releasing the trapping force, the spores might either remain adhered or separate due to Brownian motions (Fig 3d-f). This method offers a unique opportunity to study the interactions between different types of spores and between spores and vegetative cells.

**Fig 3.**
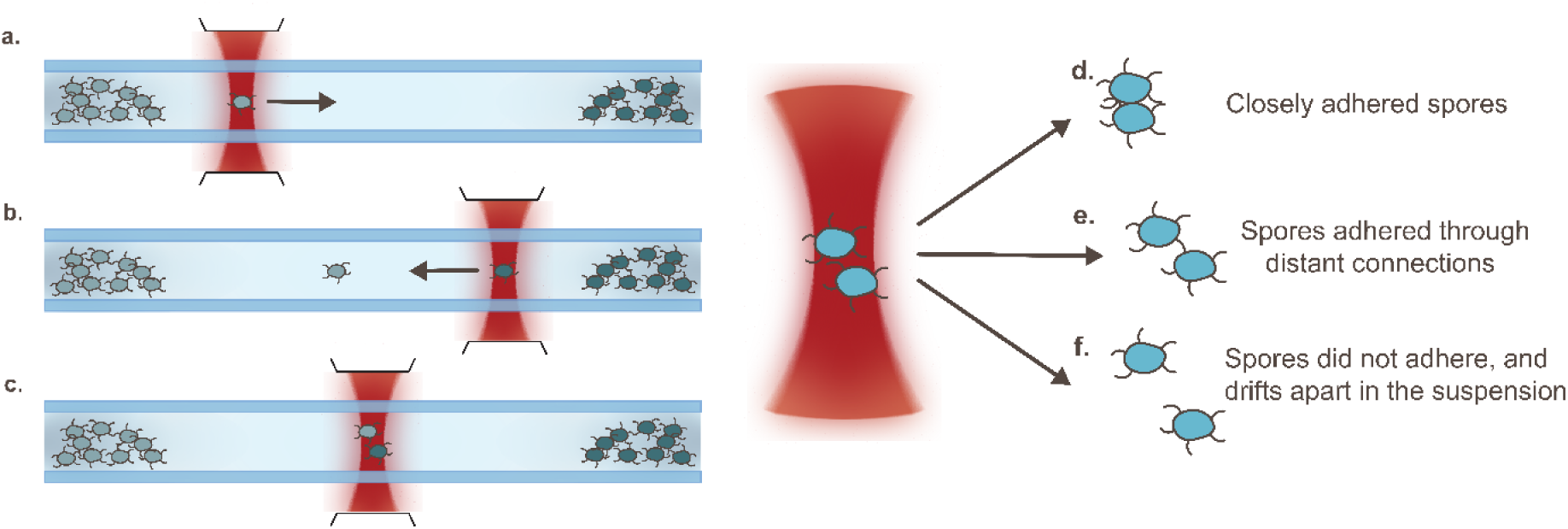
Optical tweezers catch-and-release assay. Spores of different genotypes are placed on opposite sides of a fluid channel. Spores are trapped on each side of a microfluidic channel with the OT and dragged into the center (a and b). They are held together in the optical trap (c). After release from the optical trap, the spores may either adhere to each other through close interactions (d) or maintain a more distant connection, (e) or separate and drift as a result of Brownian motion (f).

For each strain, a spore was trapped and paired with another spore of the same genotype 30 times, and the frequency of attachment – defined as instances where the two spores remained together instead of drifting apart upon release of the optical trap – was recorded (Fig 4A). WT spores remained attached in 26.7% (16/60) of the time, a frequency surpassed only by S-L+ spores, which showed a 51.6% (32/62) adherence frequency. The lowest attachment frequency was observed in the S-L+ Δ*L-bclA* spores, which displayed an even less attachment than the bald spores. Consistent with sedimentation assay results, spores lacking L-BclA (S-L+ Δ*L*-*bclA*) demonstrated significantly reduced binding compared to those with intact L-BclA. This trend toward further reduced binding in Δ*L-bclA* spores, especially when also lacking S-ENAs, is evident from the S+L+ Δ*L*-*bclA* spores’ inability to remain attached after the trapping force was removed. Altogether, the results from the catch-and-release assay correspond well with what we observed in the sedimentation assay. The strains that settled faster to the bottom of the glass tubes, were also the ones that were most likely to stick together when released from the optical trap.

**Fig 4.**
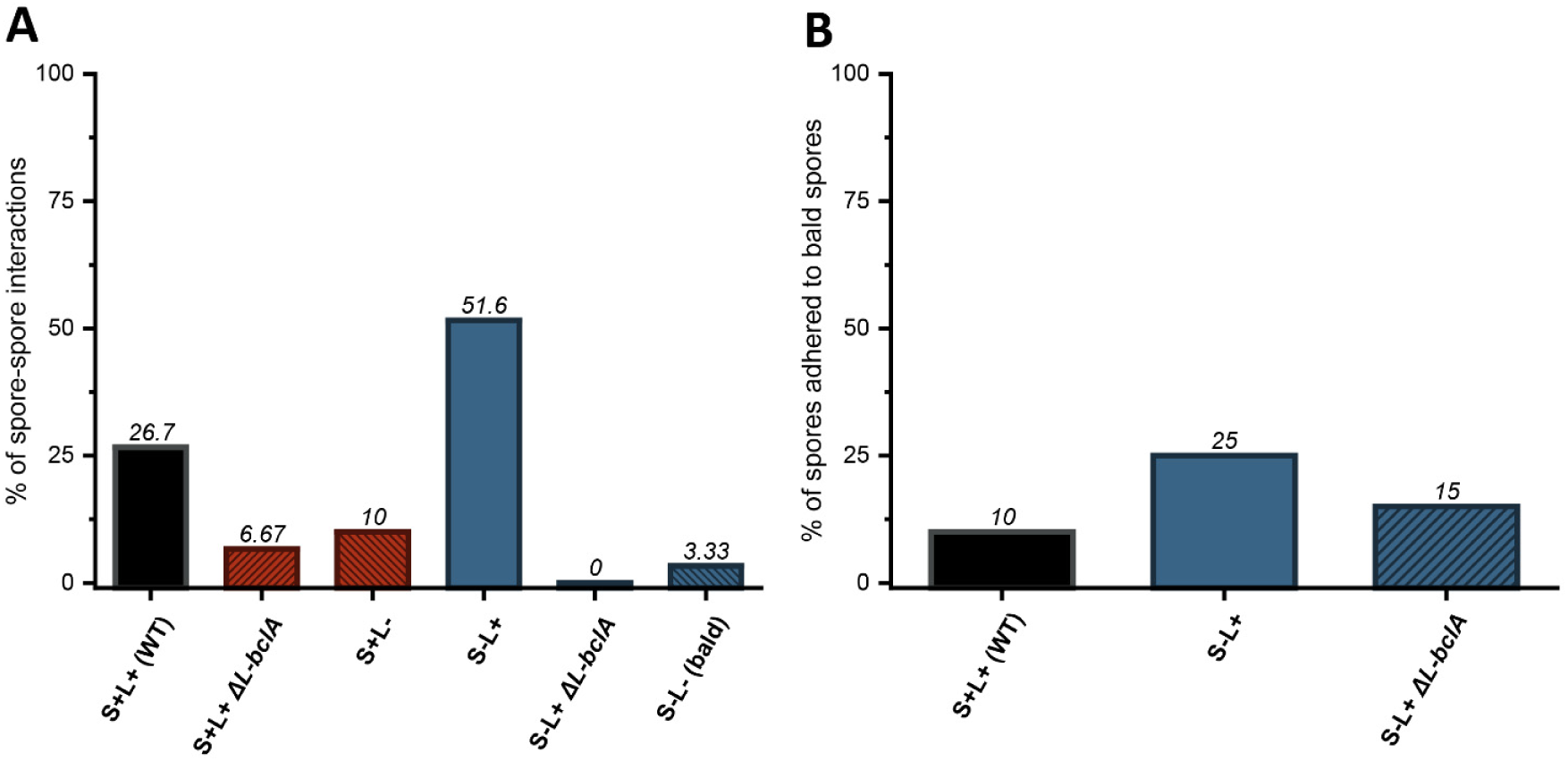
Percentage of adhered spores in water. Spores adhered to isogenic spores using the OT assay, with n = 30 interactions for all strains except S+L+ (WT) and S-L+, where n = 60, (A) and spores adhered to bald spores (B), where n = 20. Number of interaction outcomes is given in Table S3. Error bars are not included as this is an observational study.

Trapping two spores of the S++L+ strain together was notably difficult, as they consistently repelled each other, preventing the successful confinement of both within the trap. Therefore, the data represents 30 attempts to trap spores together rather than 30 actual interactions. Out of these attempts, trapping was successful in only six cases, and among these, binding was observed in only three instances. Interestingly, the binding that occurred was more distant than the close contact seen with WT or S+L-spores; typically, when one spore was trapped, the other was either pushed away or drawn closer, as seen in Figs 5A and B. The average center-to-center distance between the S++L+ spores was significantly larger at 5.8 ± 0.8 µm, nearly three times the distance of aggregated WT spores, which was 1.9 ± 0.3 µm. In the other 24 trapping attempts, only one spore remained in the trap at any given time.

**Fig 5.**
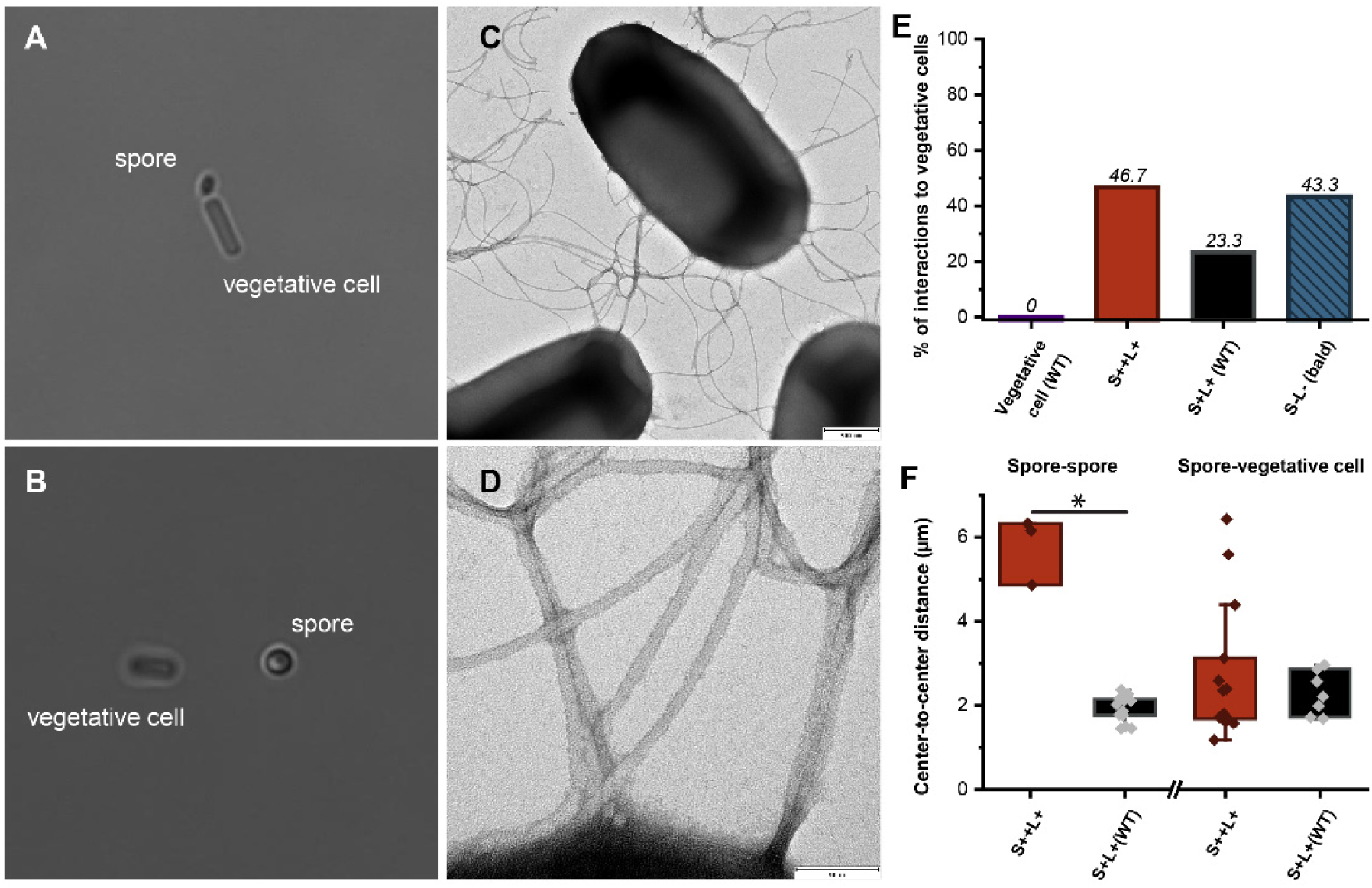
Spore-cell aggregation. Close aggregation of a WT (S+L+) spore and a vegetative cell (WT) (A) from the catch-and-release assay. Distantly connected S++L+ spore with a vegetative cell (WT) after the catch-and-release assay (B). Here the spore is picked up by the OT and the vegetative cell followed along. The surface of the vegetative cells is also decorated with flagella-like appendages (average width of 15.3 ± 2.5 nm) with distinct morphology from the endospore appendages (C, D). The bar chart shows the percentage of spores adhered to vegetative cells (E), and the boxplot gives the interaction distances (µm) between spores and between spores and vegetative cells (F). The asterisk symbolizes a significant difference (p<0.05). Average distances are given in Table S4.

**Fig 6.**
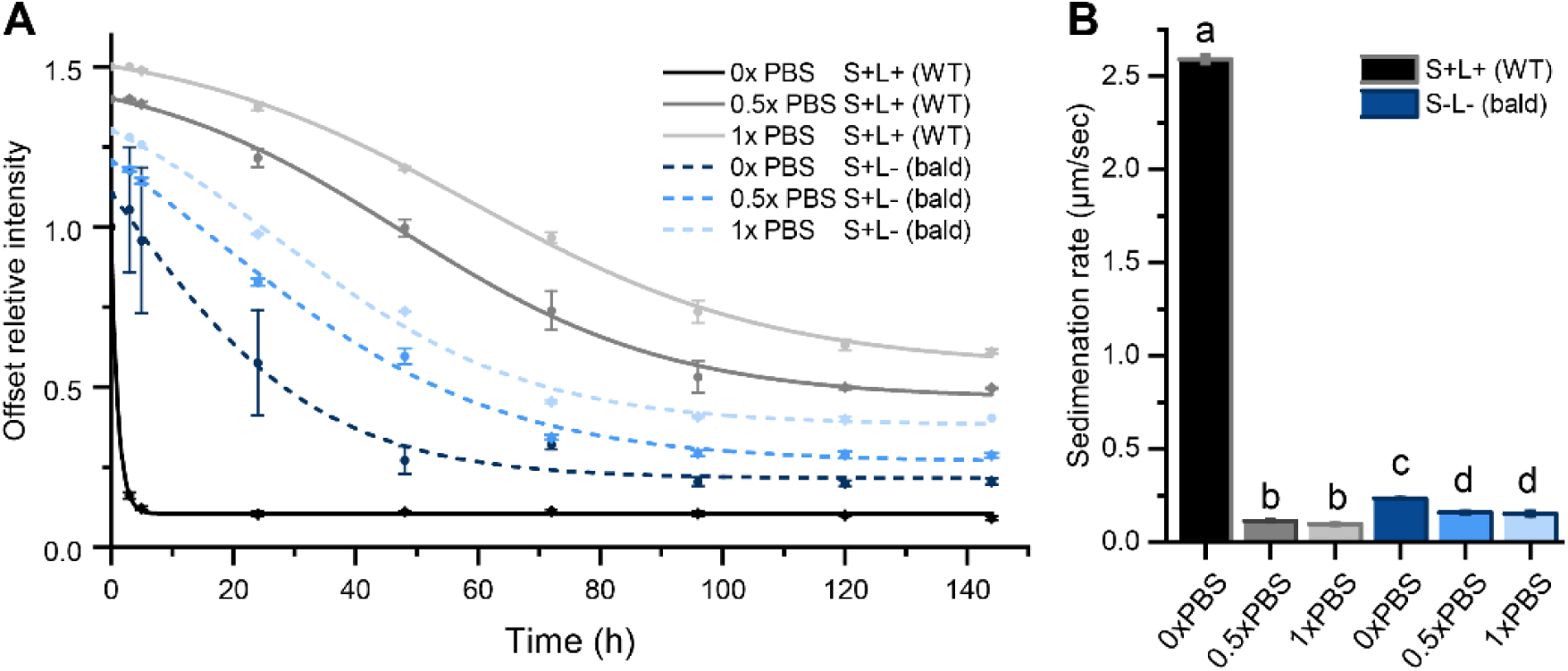
Sedimentation patterns of spores in water or PBS. Sedimentation of spores in water or PBS over time for WT (S+L+) and the ENA depletion mutant (S-L-) of *B. paranthracis* NVH 0075/95 with fitted Boltzmann curves (A), and calculated sedimentation rate in µm/s (B). Bars that share a letter are not significantly different, as indicated by Tukey’s grouping (p<0.05). For overlaying sedimentation patterns, see Figure S1 and numerical averages for sedimentation rates are given in Table S6.

Next, to test if binding requires the presence of ENAs on both interacting spores, we again performed catch-and-release experiments. In this test, we assessed the binding capabilities of WT S+L+, S-L+ and S-L+ ΔL-*bclA* spores when paired with S-L-(bald) spores, see Fig 4B. We observed that having S+L+ ENAs on only one of the spores resulted in a slightly reduced binding frequency. However, we noted that the S-L+ spores exhibited a marginally higher binding frequency than WT spores (5/20 and 2/20, respectively). In contrast to the lack of binding seen in the experiment where interaction between genetically identical spores was tested, successful binding was observed between S-L+ ΔL-*bclA* and bald spores in three out of 20 captures.

### Spore-to-vegetative cell aggregation is not dependent on ENAs

Since *B. cereus* spp. communities comprise both spores and vegetative cells, we next tested whether spores can attach to their vegetative counterpart and the potential role of ENAs in such interaction. By using the catch-and release assay, we detected frequent binding between vegetative cells *B. paranthrasis* NVH 0075/95 and WT spores (23.3%), spores that express longer S-ENAs (S++L+ (46.7%)) and bald spores (43.3%) (Fig 5E). Unlike spore-spore binding, in most cases the interaction between spores and vegetative cells occurred in much closer proximity, and the spores were observed to mainly adhere to the poles of the vegetative cells (Fig 5A). We detected two instances of more distant connections between a spore and a vegetative cell for the S++L+ mutant. In these cases, when the spore was manipulated with the OT, the vegetative cell followed along (as illustrated in Fig 5B). The interaction distance (center-to-center) between the vegetative cell and the spore was 5.6 ± 2.5 µm and 6.4 ± 0.8 µm for the two instances where this occurred. Finally, we also tested interactions between vegetative cells, but no aggregation was observed.

### PBS disrupts the binding of isogenic spores

To investigate how the aggregation behavior is affected by environmental conditions, we repeated the sedimentation experiments of WT (S+L+) spores and bald (S-L-) spores suspended in different dilutions of PBS (Fig 6). We observed that PBS in the suspension drastically reduced the sedimentation rate of the WT strain by about 2-orders of magnitude from 2.59 ± 0.01 µm/sec to 0.10 ± 0.01 µm/sec. Similarly, a significant reduction was observed in the sedimentation rate of the bald spores, with a reduction of 35% from 0.23 ± 0.01 µm/sec to 0.15 ± 0.01 µm/sec. We observed no difference in the size of the pellets between WT and bald spores after six days of settling with PBS in the suspension, Figure S5. However, the pellet of the WT strain suspended in water was significantly larger than when PBS was added to the suspension. This difference in pellet size between water and PBS was not observed in the sedimentation of bald spores.

To test if we could detect similar interference in aggregation at the single spore level using the catch-and-release assay, we examined the behavior of WT, and S-L+ spores that were most prone to aggregation in water. As shown in Fig 7, almost no interaction was observed when the spores were suspended in 1x PBS. Already at a ten-fold dilution of PBS (total ionic strength of 15.05 mM) a strong reduction in spore-spore interactions was observed. However, the addition of 0.05% of the non-ionic surfactant Tween-20 or 1% bovine serum albumin (BSA) did not result in any significant alteration in the interaction frequency between isogenic spores.

**Fig 7.**
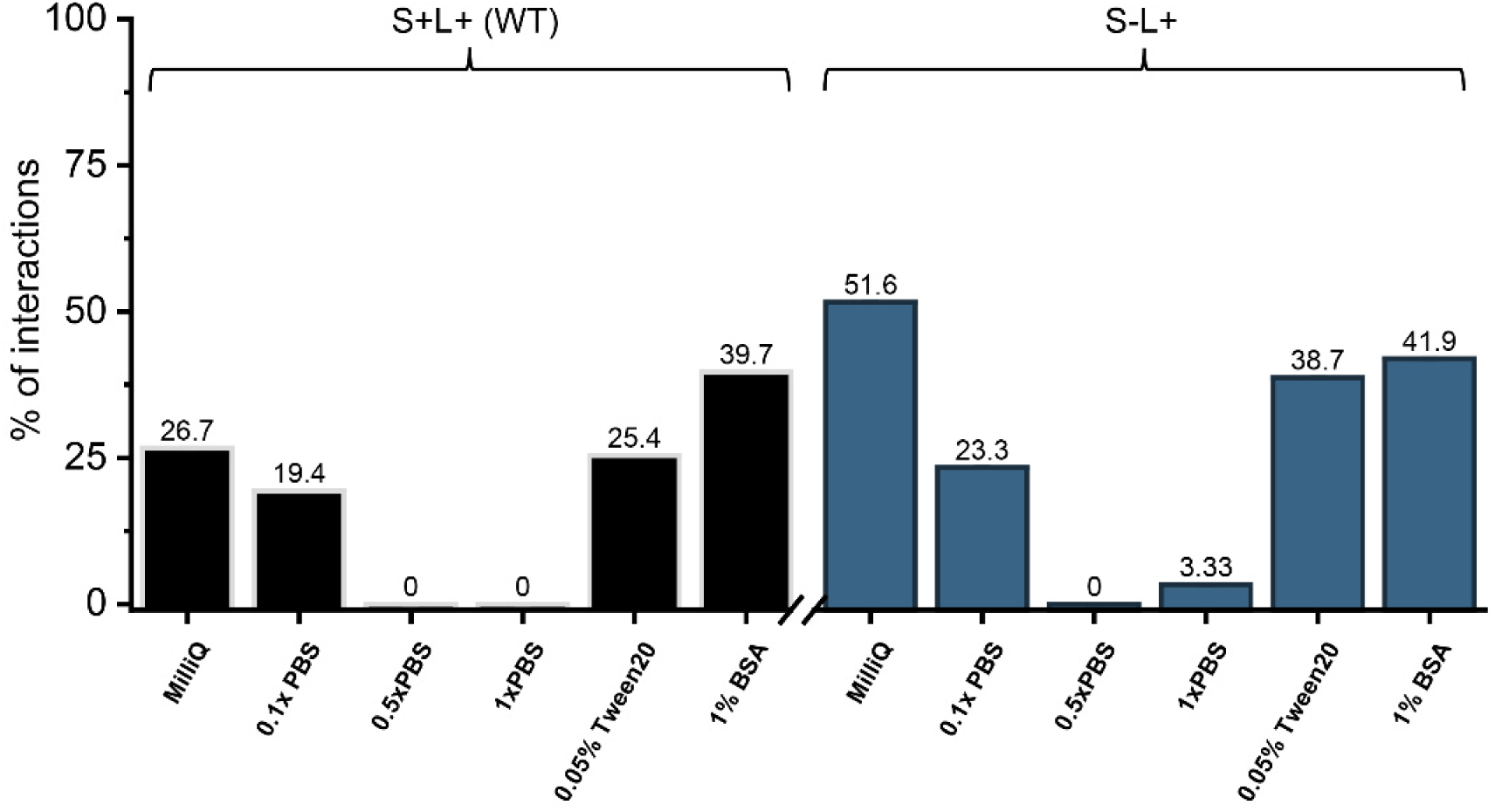
Spore interactions in water, PBS, Tween-20, and BSA solutions. The percentage of self-adherence observed during 60 and 30 interactions for WT and S-L+ strain, respectively, using the OT setup. The comparative interaction results obtained in water are the same as those shown in Fig 4A. Number of interaction outcomes is given in Table S7.

## Discussion

The present work builds upon the research conducted by Jonsmoen et al. (2023), where single-cell analyses demonstrated the role of S-ENAs in maintaining spores in a gel-like state by mediating distant spore-to-spore interactions (Jonsmoen et al., 2023). Here, by utilizing both bulk and single-cell approaches, we undertook an investigation of the role of L-and S-ENAs, as well as the tip fibrillum of the L-ENAs in the aggregative behavior of *B. paranthracis* spores, and how different factors in the surrounding liquid influences contacts between spores. We also tested whether spores adhere to vegetative cells and the role of ENAs in such interaction.

We identified two primary sedimentation patterns influenced by the ENAs. The first pattern exhibits a rapid sedimentation rate (2.40 ± 0.20 µm/sec), which depends on the presence of S-or L-ENAs. This is demonstrated by comparable sedimentation rates for S+L-(1.33 ± 0.14 µm/sec) and the S-L+ (1.23 ± 0.11 µm/sec) spores. The second pattern shows slower sedimentation rates and includes S-L-(bald) spores (0.16 ± 0.01 µm/sec), exosporium deficient spores (Δ*exsY*) (0.16 ± 0.01 µm/sec), and spores lacking both S-ENA and L-BclA (S-L+ Δ*L-bclA)*, which indicate a disruption of L-ENA-mediated spore-spore binding. Interestingly, the S+L+ Δ*L-bclA* mutant also demonstrated a significantly reduced sedimentation rate compared to the S+L-spores, in which the aggregation behavior is only supported by S-ENA. This suggests that the absence of L-BclA also influences spore aggregation mediated by S-ENA.

Noteworthy, the tip fibrilla of S-ENA closely resembles those of L-Ena, both having a diameter of 2 nm and terminating in a globular domain (Sleutel et al., 2023). Consistent with the finding that depletion of L-BclA also affects S-ENA mediated interactions between spores, TEM images comparing the S-ENA tip fibrilla of WT spores with those of S+ L+ Δ*L-bclA* spores revealed the absent fibers corresponding to the length of the L-BclA on L-ENA on S+L+ Δ*L-bclA* spores. This suggests that L-BclA may also be present on the surface of S-ENA. The lack of S-ENA mediated spore-spore adhesion resulting from the depletion of L-BclA was confirmed with the catch-and-release assay.

Despite the depletion of L-BclA, the consistent number of tip fibrilla still present on S-Enas on the S+ L+ Δ*L-bclA* suggests the presence of BclA homologs, which likely fill the void left by its absence (Sleutel et al., 2023). After identifying L-BclA as a functional component of the S-ENA-fibers, it is intriguing to consider that the genetic identity of the other tip fibrilla on S-ENAs remains unknown. In the NVH0075/95 genome, there are 12 collagen-like genes, which lack the sequence encoding the exosporium localization domain proteins (Files S8-10). It’s plausible that one or several of these homologs serve as “alternative” S-Ena tip fibrilla.

Furthermore, while the *ena3* gene cluster is only present in about 10% of the *B. cereus* s. l. genomes, almost all other genomes carry the *ena1/2* gene clusters (Pradhan et al., 2021). Assuming that all these ENA fibers exhibit tip fibrilla, it is likely that these structures are encoded by BclA homologues other than L-BclA (Sleutel et al., 2023). However, whether such homologs also facilitate clustering of spores or serve other functions requires further investigation.

In addition to the reduced sedimentation rate of the Δ*L-bclA* depleted strain (S+L+ Δ*L-bclA*) we also observed that the S++ L+ strain sediments significantly slower than the WT strain. They also showed a distinct sedimentation behavior by settling more collectively in contrast to the more diffuse sedimentation pattern observed for spores of the other strains (Fig 1B). The pellet that was formed by S++ L+ spores at the bottom of the tube, was also less dense compared to those of the other spore suspensions. The resistance to dense packing suggests that the S-ENAs present a steric hindrance preventing close encounters between spores. The S++L+ spore pellet was gradually compressed over the days the experiment continued, indicating that the steric hindrance was reduced over time. In Jonsmoen et al., 2023, we demonstrated that spores expressing S-ENA exhibit a gel-like state when suspended in water i.e., a viscosity that indicates relaxed interconnection between spores (Jonsmoen et al., 2023). The gel-like behavior was not observed for ENA-depleted bald spores, which moved independently of each other when suspended in water. The concept that S-ENAs act as a physical barrier, hindering close encounters between spores, is supported by the difficulty of capturing two S++L+ spores simultaneously in the optical tweezer trap.

We also demonstrate that the composition of the surrounding medium influences the binding between spores. During sedimentation of spores, we observed a significant reduction in the sedimentation rate of both WT (S+L+) and bald (S-L-) spores when PBS was added to the surrounding medium. A similar reduction of binding was observed during the catch-and-release assay where we tested WT (S+L+) spores and spores of the strain only expressing L-ENAs (S-L+), which both showed the highest tendency to bind in water. When PBS was added to the spore suspension, the density and ionic strength of the suspension media increases, affecting its resistance towards sedimentation and the level of free ions that may influence protein interactions. In biological research, PBS is commonly used to preserve biological function by being isotonic and keeping a physiological pH (7.4). This is, however, not necessarily a relevant condition for all bacteria, as the salinity differs between different environmental habitats. Therefore, we tested how lower salt concentrations influenced spore-binding in the catch-and-release assay. Already, with a salt concentration of 15 mM in the surrounding liquid medium (0.1x PBS), we noted a decrease in spore aggregation at the single spore level, while at higher concentrations, the spores failed to aggregate. The decreased binding observed at increased ionic strength suggests that the interactions observed are specific and charge dependent.

Examining the frequencies of spore aggregation in different liquid suspensions provides valuable insights into the mechanisms driving spore-spore adhesion. It is, however, important to acknowledge the role played by the non-specific physicochemical surface properties of bacteria in autoaggregation (Trunk et al., 2018). When the non-ionic surfactant Tween-20 was added to the suspension, we observed no decrease in binding affinity, indicating that non-specific hydrophobic interactions were not the driving force behind ENA-mediated spore-spore interactions. Similarly, the presence of the protein blocking agent BSA did not inhibit the spores from binding.

Investigations into spore interaction in different conditions are important for increasing our understanding of their collective behavior and their interactions with their environment. We speculate that the aggregation properties of the spores may contribute to their persistence in the upper layers of the soil, where nutrient and organic matter concentrations are higher compared to deeper layers. In these surface layers, spores are more accessible to grazing animals (Hugh-Jones & Blackburn, 2009; Vissers et al., 2007). From the soil, spores enter our food production systems through contact with animals and raw food materials (Carlin, 2011; Christiansson et al., 1999; Heyndrickx, 2011). In the food industry, spores pose a significant risk as they survive heat treatment and other disinfection methods that are implemented to ensure food safety and high quality of food products (ENEROTH et al., 2001; Heyndrickx, 2011). Surviving spores may germinate, grow and form biofilms, a recurrent problem in food production facilities (Kumari & Sarkar, 2016; Shaheen et al., 2010b). Biofilms of *B. cereus* consist of substantial amounts of spores (Wijman et al., 2007). As the biofilms mature, the levels of spores fluctuate (Ryu & Beuchat, 2005) due to shedding or germination, with remnants of spores becoming a part of the biofilm matrix. Spore appendages are highly resilient to proteinases and other degrading enzymes (Pradhan et al., 2021) and may accumulate in biofilm over time. The tensile stiff S-ENAs may consequently become an important constituent of the biofilm matrix, contributing to its structure and rigidity. This hypothesis is further supported by the ability of ENAs to aggregate spores, as well as to create a gel-like state of interconnected spores in a suspension. The steric resistance of the S-ENA fiber might also prevent too close encounters between neighboring spores and cells which may improve the flow of nutrients through the biofilm. Further, the greater hydrodynamic diameter attributed to the presence of appendages on the spores (Jonsmoen et al., 2023) may ease spore detachment from the outer layer of the biofilm and help spread spores into the environment. Even so, we observed no effect of ENAs in binding of spores to vegetative cells.

The ability of bacteria to form aggregates is associated with pathogenesis. The effect is passive, but beneficial as it protects the bacteria from microbicidal agents, harsh environmental conditions, and host defenses (Trunk et al., 2018). The prolonged persistence enhances the chance of survival and for successful colonization of the host. In the case of *B. cereus* infection, where a high infectious dose of 10^5^ CFU/mL is required (Granum & Lund, 2006), the aggregation of spores might play a pivotal role. By sticking together, the spores enhance their likelihood of reaching the host gut, where they germinate and initiate infection. While the germination behavior of spores in aggregates has not been investigated, it is known that the release of CaDPA triggers germination of surrounding spores (Setlow, 2014), suggesting a well-coordinated germination behavior in a spore population. Yet, in a population of spores, a small fraction is considered to be hyper dormant (Ghosh & Setlow, 2009), a bet-hedging strategy where some spores resist cues that typically trigger germination, potentially providing an additional evolutionary advantage for aggregating spores.

## Conclusion

This study aimed to explore the role of ENAs in spore-to-spore and spore-to-vegetative cells aggregation. Our findings, obtained through ensemble sedimentation studies and single cell techniques, reveal the involvement of both S-and L-ENAs in the aggregation of *B. paranthracis* NVH 0075/95 spores and highlight the role of the L-BclA tip fibrilla as a crucial functional element of both ENA types (Fig 8). While we observe that the ENAs are not important for binding of spores-to-vegetive cells, we recognize the potential benefits of their rigidity and stiffness in enhancing the rigidity and functionality of a biofilm matrix. Furthermore, our results demonstrate that the aggregation mechanism is sensitive to the surrounding environment of the spores, being disrupted by higher ionic strength in the suspension while remaining unaffected by surfactants. This suggests that spore interactions are charge dependent and affected by environmental changes.

**Fig 8.**
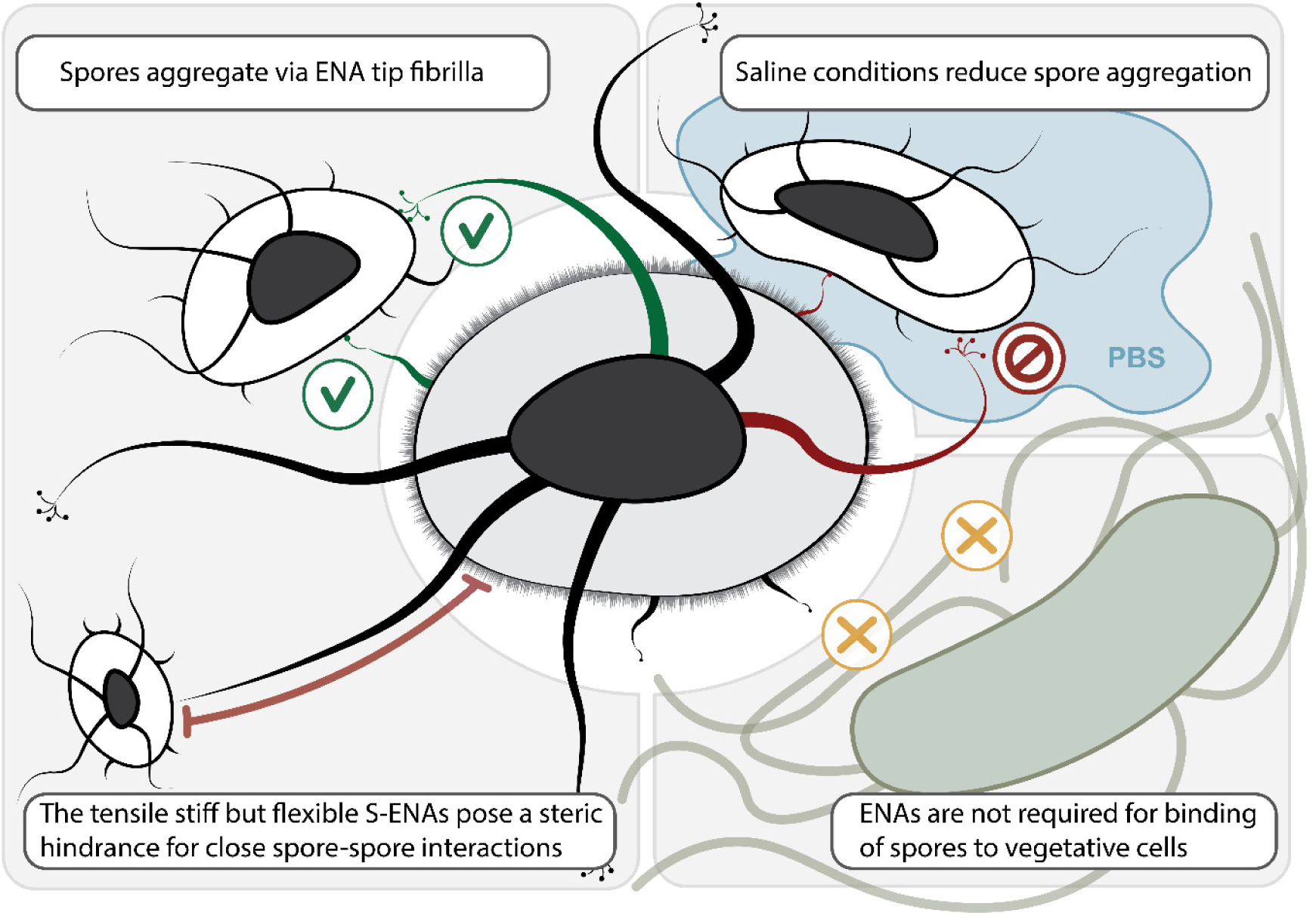
Graphical conclusion. Spore aggregation is mediated through ENAs and likely through the ENA tip-fibrilla, particularly important are the L-BclA. S-ENAs provide a steric hindrance for close interactions. The ENAs are not involved in adhesion to vegetative cells, and the spore-spore aggregation is reduced by salt in the surrounding media.

## Acknowledgements

The authors acknowledge the facilities and technical assistance of the NMBU Imaging Center and PhD student Jennie Ann Allred for TEM image of vegetative *B. cereus* NVH 0075/95 cells. A special thanks to Yohannes Beyene Mekonnen for technical assistance and Kristin (Tina) O’Sullivan for constructing mutants.

## Data Availability Statement

All data are included within the article or as supporting information.

## Supporting information captions

**S1 Figure. Sedimentation profiles of spores in water (A) and PBS (B).**

**S2 Table. Sedimentation rates (Average ± SD) and pellet size in MilliQ-water. S3 Table. Number of close interactions for catch-and-release assay in Milli-Q. S4 Table. Center-to-center distances for catch-and-release tests.**

**S5 Figure. Pellet hight for spores sedimenting in PBS and water.** Bars that share a letter are not significantly different, as indicated by Tukey’s grouping (p<0.05).

**S6 Table. Sedimentation rate (Average ± SD) in PBS.**

**S7 Table. Distribution and number of close interactions for catch-and-release assay in different liquid media.**

**S8 Text. Description of HMMER search for collagen-like homologes in NVH0075/95 S9 File. Bacillus_paranthracis_NM_GCF_027945115.1_hmmer.log**

**S10 File. NM_hits_collagen_leader_sequence_hmm.log**

## References

Andersson, M., Czerwinski, F., & Oddershede, L. B. (2011). Optimizing active and passive calibration of optical tweezers. Journal of Optics, 13(4), 044020. 10.1088/2040-8978/13/4/044020

Bieber, D., Ramer, S. W., Wu, C.-Y., Murray, W. J., Tobe, T., Fernandez, R., & Schoolnik, G. K. (1998). Type IV Pili, Transient Bacterial Aggregates, and Virulence of Enteropathogenic *Escherichia coli*. Science, 280(5372), 2114–2118. 10.1126/science.280.5372.2114

Boyer, R. R., Sumner, S. S., Williams, R. C., Pierson, M. D., Popham, D. L., & Kniel, K. E. (2007). Influence of Curli Expression by Escherichia coli O157:H7 on the Cell’s Overall Hydrophobicity, Charge, and Ability To Attach to Lettuce. Journal of Food Protection, 70(6), 1339–1345. 10.4315/0362-028X-70.6.1339

Burel, C., Dreyfus, R., & Purevdorj-Gage, L. (2021). Physical mechanisms driving the reversible aggregation of Staphylococcus aureus and response to antimicrobials. Scientific Reports, 11(1), 15048. 10.1038/s41598-021-94457-1

Carlin, F. (2011). Origin of bacterial spores contaminating foods. Food Microbiology, 28(2), 177–182. 10.1016/j.fm.2010.07.008

Christiansson, A., Bertilsson, J., & Svensson, B. (1999). Bacillus cereus Spores in Raw Milk: Factors Affecting the Contamination of Milk During the Grazing Period. Journal of Dairy Science, 82(2), 305–314. 10.3168/jds.S0022-0302(99)75237-9

Demirdjian, S., Sanchez, H., Hopkins, D., & Berwin, B. (2019). Motility-Independent Formation of Antibiotic-Tolerant Pseudomonas aeruginosa Aggregates. Applied and Environmental Microbiology, 85(14). 10.1128/AEM.00844-19

Dogsa, I., Kostanjšek, R., & Stopar, D. (2023). eDNA Provides a Scaffold for Autoaggregation of B. subtilis in Bacterioplankton Suspension. Microorganisms, 11(2). 10.3390/MICROORGANISMS11020332

Eneroth, Å., Svensson, B., Molin, G., & Christiansson, A. (2001). Contamination of pasteurized milk by *Bacillus cereus* in the filling machine. Journal of Dairy Research, 68(2), 189–196. 10.1017/S002202990100485X

Faille, C., Bénézech, T., Midelet-Bourdin, G., Lequette, Y., Clarisse, M., Ronse, G., Ronse, A., & Slomianny, C. (2014). Sporulation of Bacillus spp. within biofilms: A potential source of contamination in food processing environments. Food Microbiology, 40, 64–74. 10.1016/j.fm.2013.12.004

Faille, C., Bihi, I., Ronse, A., Ronse, G., Baudoin, M., & Zoueshtiagh, F. (2016). Increased resistance to detachment of adherent microspheres and Bacillus spores subjected to a drying step. Colloids and Surfaces B: Biointerfaces, 143, 293–300. 10.1016/j.colsurfb.2016.03.041

Faris, A., Lindahl, M., Ljungh, Å., Old, D. C., & Wadström, T. (1983). Autoaggregating *Yersinia enterocolitica* express surface fimbriae with high surface hydrophobicity. Journal of Applied Bacteriology, 55(1), 97–100. 10.1111/j.1365-2672.1983.tb02652.x

Ghosh, S., & Setlow, P. (2009). Isolation and Characterization of Superdormant Spores of *Bacillus* Species. Journal of Bacteriology, 191(6), 1787–1797. 10.1128/JB.01668-08

Granum, P. E., & Lund, T. (2006). Bacillus cereus and its food poisoning toxins. FEMS Microbiology Letters, 157(2), 223–228. 10.1111/j.1574-6968.1997.tb12776.x

Heyndrickx, M. (2011). The Importance of Endospore-Forming Bacteria Originating from Soil for Contamination of Industrial Food Processing. Applied and Environmental Soil Science, 2011, 1–11. 10.1155/2011/561975

Hugh-Jones, M., & Blackburn, J. (2009). The ecology of Bacillus anthracis. Molecular Aspects of Medicine, 30(6), 356–367. 10.1016/j.mam.2009.08.003

Jonsmoen, U. L., Malyshev, D., Öberg, R., Dahlberg, T., Aspholm, M. E., & Andersson, M. (2023). Endospore pili: Flexible, stiff, and sticky nanofibers. Biophysical Journal, 122(13), 2696–2706. 10.1016/j.bpj.2023.05.024

Kragh, K. N., Hutchison, J. B., Melaugh, G., Rodesney, C., Roberts, A. E. L., Irie, Y., Jensen, P. Ø., Diggle, S. P., Allen, R. J., Gordon, V., & Bjarnsholt, T. (2016). Role of Multicellular Aggregates in Biofilm Formation. MBio, 7(2). 10.1128/mBio.00237-16

Kumari, S., & Sarkar, P. K. (2016). Bacillus cereus hazard and control in industrial dairy processing environment. Food Control, 69, 20–29. 10.1016/j.foodcont.2016.04.012

Lund, T., & Granum, P. E. (1996). Characterisation of a non-haemolytic enterotoxin complex from *Bacillus cereus* isolated after a foodborne outbreak. FEMS Microbiology Letters, 141(2–3), 151–156. 10.1111/j.1574-6968.1996.tb08377.x

Malyshev, D., Öberg, R., Dahlberg, T., Wiklund, K., Landström, L., Andersson, P. O., & Andersson, M. (2022). Laser induced degradation of bacterial spores during micro-Raman spectroscopy. Spectrochimica Acta Part A: Molecular and Biomolecular Spectroscopy, 265, 120381. 10.1016/J.SAA.2021.120381

Nwoko, E. Q. A., & Okeke, I. N. (2021). Bacteria autoaggregation: how and why bacteria stick together. Biochemical Society Transactions, 49(3), 1147–1157. 10.1042/BST20200718

Peng, J.-S., Tsai, W.-C., & Chou, C.-C. (2001). Surface characteristics of Bacillus cereus and its adhesion to stainless steel. International Journal of Food Microbiology, 65(1–2), 105–111. 10.1016/S0168-1605(00)00517-1

Pradhan, B., Liedtke, J., Sleutel, M., Lindbäck, T., Zegeye, E. D., ÓSullivan, K., Llarena, A., Brynildsrud, O., Aspholm, M., & Remaut, H. (2021). Endospore Appendages: a novel pilus superfamily from the endospores of pathogenic Bacilli. The EMBO Journal, 40(17). 10.15252/embj.2020106887

Rahnama, H., Azari, R., Yousefi, M. H., Berizi, E., Mazloomi, S. M., Hosseinzadeh, S., Derakhshan, Z., Ferrante, M., & Conti, G. O. (2023). A systematic review and meta-analysis of the prevalence of Bacillus cereus in foods. Food Control, 143, 109250. 10.1016/j.foodcont.2022.109250

Rönner, U., Husmark, U., & Henriksson, A. (1990). Adhesion of bacillus spores in relation to hydrophobicity. Journal of Applied Bacteriology, 69(4), 550–556. 10.1111/j.1365-2672.1990.tb01547.x

Rueden, C. T., Schindelin, J., Hiner, M. C., DeZonia, B. E., Walter, A. E., Arena, E. T., & Eliceiri, K. W. (2017). ImageJ2: ImageJ for the next generation of scientific image data. BMC Bioinformatics, 18(1), 529. 10.1186/s12859-017-1934-z

Ryu, J.-H., & Beuchat, L. R. (2005). Biofilm Formation and Sporulation by Bacillus cereus on a Stainless Steel Surface and Subsequent Resistance of Vegetative Cells and Spores to Chlorine, Chlorine Dioxide, and a Peroxyacetic Acid–Based Sanitizer. Journal of Food Protection, 68(12), 2614–2622. 10.4315/0362-028X-68.12.2614

Setlow, P. (2006). Spores of Bacillus subtilis: their resistance to and killing by radiation, heat and chemicals. Journal of Applied Microbiology, 101(3), 514–525. 10.1111/j.1365-2672.2005.02736.x

Setlow, P. (2014). Germination of Spores of Bacillus Species: What We Know and Do Not Know. Journal of Bacteriology, 196(7), 1297–1305. 10.1128/JB.01455-13

Shaheen, R., Svensson, B., Andersson, M. A., Christiansson, A., & Salkinoja-Salonen, M. (2010a). Persistence strategies of Bacillus cereus spores isolated from dairy silo tanks. Food Microbiology, 27(3), 347–355. 10.1016/j.fm.2009.11.004

Shaheen, R., Svensson, B., Andersson, M. A., Christiansson, A., & Salkinoja-Salonen, M. (2010b). Persistence strategies of Bacillus cereus spores isolated from dairy silo tanks. Food Microbiology, 27(3), 347–355. 10.1016/j.fm.2009.11.004

Simmonds, P., Mossel, B. L., Intaraphan, T., & Deeth, H. C. (2003). Heat Resistance of Bacillus Spores When Adhered to Stainless Steel and Its Relationship to Spore Hydrophobicity. Journal of Food Protection, 66(11), 2070–2075. 10.4315/0362-028X-66.11.2070

Sleutel, M., Zegeye, E. D., Llarena, A. K., Pradhan, B., Fislage, M., O’Sullivan, K., Aspholm, M., & Remaut, H. (2023). A novel class of ultra-stable endospore appendages decorated with collagen-like tip fibrillae. BioRxiv, 2023.10.23.563578. 10.1101/2023.10.23.563578

Stangner, T., Dahlberg, T., Svenmarker, P., Zakrisson, J., Wiklund, K., Oddershede, L. B., & Andersson, M. (2018). Cooke–Triplet tweezers: more compact, robust, and efficient optical tweezers. Optics Letters, 43(9), 1990. 10.1364/OL.43.001990

Sunde, E. P., Setlow, P., Hederstedt, L., & Halle, B. (2009). The physical state of water in bacterial spores. Proceedings of the National Academy of Sciences, 106(46), 19334– 19339. 10.1073/pnas.0908712106

Swick, M. C., Koehler, T. M., & Driks, A. (2016). Surviving Between Hosts: Sporulation and Transmission. Microbiology Spectrum, 4(4). 10.1128/microbiolspec.VMBF-0029-2015

Trunk, T., S. Khalil, H., & C. Leo, J. (2018). Bacterial autoaggregation. AIMS Microbiology, 4(1), 140–164. 10.3934/microbiol.2018.1.140

Vissers, M. M. M., Te Giffel, M. C., Driehuis, F., De Jong, P., & Lankveld, J. M. G. (2007). Predictive Modeling of Bacillus cereus Spores in Farm Tank Milk During Grazing and Housing Periods. Journal of Dairy Science, 90(1), 281–292. 10.3168/jds.S0022-0302(07)72629-2

Wijman, J. G. E., de Leeuw, P. P. L. A., Moezelaar, R., Zwietering, M. H., & Abee, T. (2007). Air-Liquid Interface Biofilms of *Bacillus cereus*: Formation, Sporulation, and Dispersion. Applied and Environmental Microbiology, 73(5), 1481–1488. 10.1128/AEM.01781-06

Yang, S., Wang, Y., Ren, F., Wang, X., Zhang, W., Pei, X., & Dong, Q. (2023). The Sources of *Bacillus cereus* Contamination and their Association with Cereulide Production in Dairy and Cooked Rice Processing Lines. Food Quality and Safety, 7. 10.1093/fqsafe/fyad023

Zegeye, E. D., Pradhan, B., Llarena, A.-K., & Aspholm, M. (2021). Enigmatic Pilus-Like Endospore Appendages of Bacillus cereus Group Species. International Journal of Molecular Sciences, 22(22), 12367. 10.3390/ijms222212367

