## Supporting Information for "The role of endospore appendages in spore-spore contacts in pathogenic bacilli"

Short title: ENA-dependent endospore aggregation

Unni Lise Jonsmoen[1], Dmitry Malyshev[2], Mike Sleutel[3,4], Elise Egeli Kristensen[1], Ephrem Debebe Zegeye[1], Han Remaut[3,4], Magnus Andersson[2,5*] and Marina Elisabeth Aspholm[1*].

[1] Department of Paraclinical Sciences, Faculty of Veterinary Medicine, Norwegian University of Life Sciences (NMBU), Ås, Norway

[2] Department of Physics, Umeå University, Umeå, Sweden

[3] Structural Biology Brussels, Vrije Universiteit Brussel, Brussels, Belgium

[4] Structural and Molecular Microbiology, Structural Biology Research Center, VIB, Brussels, Belgium

[5] Umeå Centre for Microbial Research (UCMR)

* Corresponding authors

**E-mail addresses of authors:** Jonsmoen:, Malyshev:, Sleutel: , Kristensen:, Zegeye:, Remaut:, Andersson:, and Aspholm::

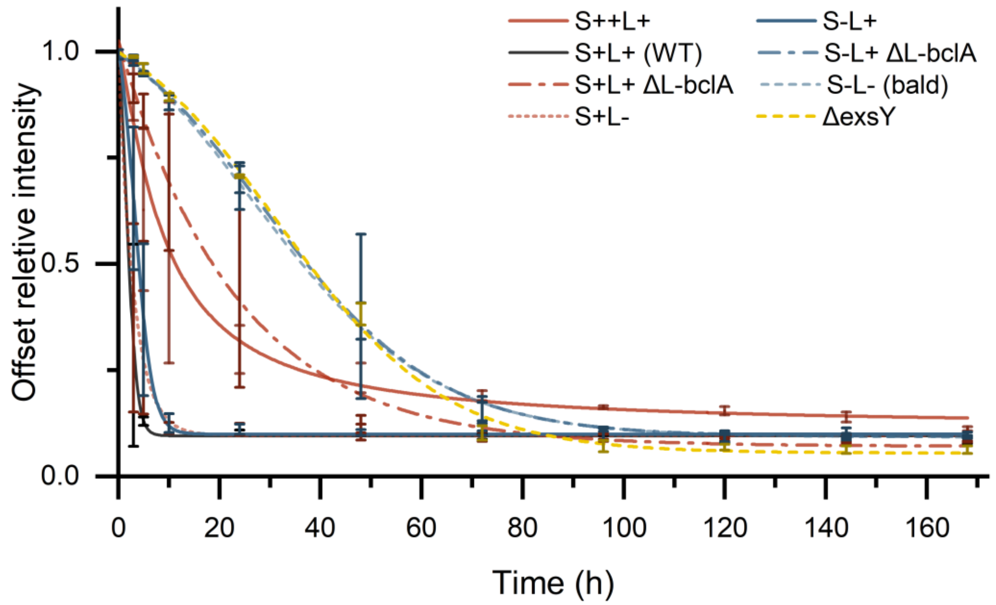


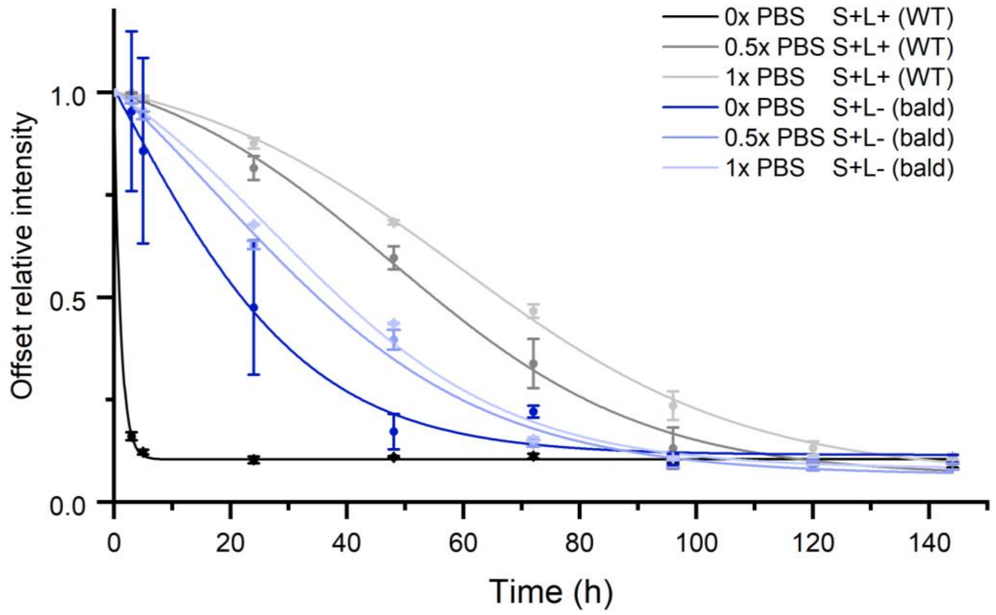


**S1 Figure. Sedimentation profiles of spores in water (A) and PBS (B).**

**S2 Table. Sedimentation rates (Average ± SD) and pellet size in MilliQ-water.**

| Strains | Sedimentation rate (μm/sec) | Pellet size (mm) |
|---|---|---|
| S++L+ | 0.42 ± 0.08 | 3.6 ± 0.1 |
| S+L+ (WT) | 2.40 ± 0.20 | 2.2 ± 0.1 |
| S+L+ Δ*L-bclA* | 0.31 ± 0.13 | 1.4 ± 0.1 |
| S+L- | 1.33 ± 0.14 | 1.6 ± 0.7 |
| S-L+ | 1.23 ± 0.11 | 2.1 ± 0.1 |
| S-L+ Δ*L-bclA* | 0.15 ± 0.01 | 1.7 ± 0.1 |
| S-L- (Bald) | 0.16 ± 0.01 | 1.6 ± 0.1 |
| Δ*exsY* | 0.16 ± 0.01 | 1.0 ± 0.1 |

**S3 Table. Number of close interactions for catch-and-release assay in Milli-Q.**

| Spore-to-spore | Yes | No | No. Of interactions | % |
|---|---|---|---|---|
| WT | 16 | 44 | 60 | 26.7 |
| Bald | 1 | 29 | 30 | 3.3 |
| S-L+ | 32 | 30 | 62 | 51.6 |
| S-L+ Δ*L-bclA* | 0 | 30 | 30 | 0 |
| S++L+ | | | | |
| S+L- | 3 | 27 | 30 | 10.0 |
| S+L+ Δ*L-bclA* | 2 | 28 | 30 | 6.7 |
| **Spore-to-veg** | **Yes** | **No** | **No. Of interactions** | **%** |

| | | | | |
|---|---|---|---|---|
| Vegetative cell | 0 | 30 | 30 | 0 |
| WT | 7 | 23 | 30 | 23.3 |
| Bald | 13 | 17 | 30 | 43.3 |
| S++L+ | 14 | 16 | 30 | 46.7 |
| **Spore-to-bald spore** | **Yes** | **No** | **No. Of interactions** | **%** |
| WT | 2 | 18 | 20 | 10 |
| S-L+ | 5 | 15 | 20 | 25 |
| S-L+ Δ*L-bclA* | 3 | 17 | 20 | 15 |

**S4 Table. Center-to-center distances for catch-and-release tests.**

| Center-to-center distance | Strain | N | Average distance (μm) |
|---|---|---|---|
| Spore-to-spore | S++L+ | 3 | 5.78 ± 0.80 |
| | S+L+ (WT) | 13 | 1.93 ± 0.32 |
| Spore-to-vegetative cell | S++L+ | 13 | 2.80 ± 1.66 |
| | S+L+ (WT) | 7 | 2.29 ± 0.53 |

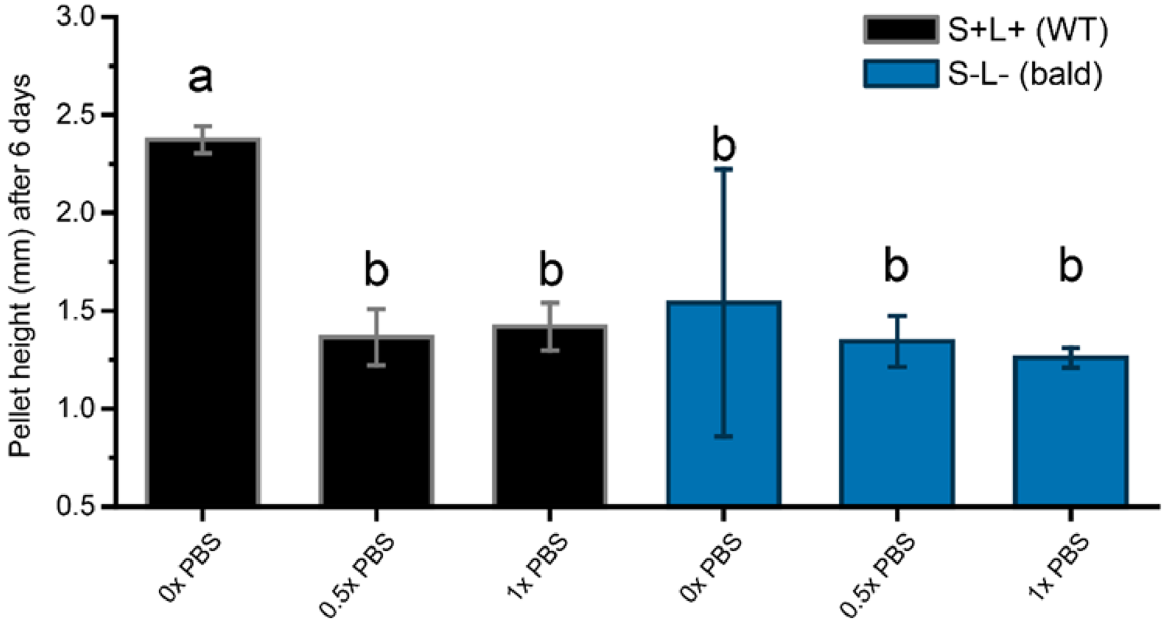


**S5 Figure. Pellet hight for spores sedimenting in PBS and water.** Bars that share a letter are not significantly different, as indicated by Tukey's grouping ($p<0.05$).

**S6 Table. Sedimentation rate (Average ± SD) in PBS.**

| **Strains** | **PBS Concentration** | **Sedimentation rate (µm/sec)** | **Pellet size (mm)** |
|---|---|---|---|
| S+L+ (WT) | 0x PBS | 2.59 ± 0.01 | 2.4 ± 0.1 |
| | 0.5x PBS | 0.11 ± 0.01 | 1.4 ± 0.1 |
| | 1x PBS | 0.10 ± 0.01 | 1.4 ± 0.1 |
| S-L- (bald) | 0x PBS | 0.23 ± 0.01 | 1.5 ± 0.4 |
| | 0.5x PBS | 0.16 ± 0.02 | 1.3 ± 0.1 |
| | 1x PBS | 0.15 ± 0.01 | 1.3 ± 0.1 |

**S7 Table. Distribution and number of close interactions for catch-and-release assay in different liquid media.**

| **Milli-Q** | **Yes** | **No** | **No. Of interactions** | **%** |
|---|---|---|---|---|
| S+L+ (WT) | 16 | 44 | 60 | 26.7 |
| S-L+ | 32 | 30 | 62 | 51.6 |
| **0.1x PBS** | **Yes** | **No** | **No. Of interactions** | **%** |
| S+L+ (WT) | 12 | 50 | 62 | 19.4 |
| S-L+ | 7 | 23 | 30 | 23.3 |
| **0.5x PBS** | **Yes** | **No** | **No. Of interactions** | **%** |
| S+L+ (WT) | 0 | 61 | 61 | 0 |
| S-L+ | 0 | 30 | 30 | 0 |
| **1x PBS** | **Yes** | **No** | **No. Of interactions** | **%** |
| S+L+ (WT) | 0 | 60 | 60 | 0 |
| S-L+ | 1 | 29 | 30 | 3.3 |
| **0.05% Tween20** | **Yes** | **No** | **No. Of interactions** | **%** |
| S+L+ (WT) | 16 | 47 | 63 | 25.4 |
| S-L+ | 12 | 19 | 31 | 38.7 |
| **1% BSA** | **Yes** | **No** | **No. Of interactions** | **%** |
| S+L+ (WT) | 25 | 38 | 63 | 39.7 |
| S-L+ | 13 | 18 | 31 | 41.9 |

**S8 Text. Description of HMMER search for collagen-like homologues in NVH0075/95**

We used the *Bacillus paranthracis* genome (RefSeq: GCF_027945115.1) to conduct a HMMER search of collagen-like protein homologues in the genome. For this, we used a custom HMM file based on the MSA of collagen-like sequences. We got 15 hits, see the log file S10. Using the NCBI_fam hmm model TIGR03720, we identified that three of these 15 hits contained exosporium leader sequences (S11 file). This leaves 12 tip fribrilla collagen-like candidates for this *B. paranthracis* strain based solely on HMM searches.

**S9 File. Bacillus_paranthracis_NM_GCF_027945115.1_hmmer.log**

**S10 File. NM_hits_collagen_leader_sequence_hmm.log**
